# Muscle-driven hand simulations emphasize the critical role of the extensor mechanism

**DOI:** 10.64898/2026.05.11.723556

**Authors:** Maximilian Carvajal, Pouyan Firouzabadi, Fabio Rizzoglio, Vikram Darbhe, R. James Cotton, Lee E. Miller, Wendy M. Murray

## Abstract

Biomechanical simulations of complex hand motions remain scarce, due to challenges that span computation and data acquisition. Using a computer vision-based motion capture approach, a 23-degree of freedom musculoskeletal model, and direct collocation optimization, we performed muscle-driven simulations to track hand kinematics from 7 participants performing American Sign Language gestures. While proximal joints were tracked accurately, interphalangeal joint tracking was significantly worse, with a consistent flexion bias. Modifications to finger extensor muscle paths that incorporated the dual-inserting nature of the extensors improved accuracy, suggesting better representation of extensor force distribution across distal joints may be necessary for accurate hand simulations.

## Introduction

The complex anatomy and sophisticated neural control of the human hand enable an extraordinary range of function that is fundamental to human capability. Loss of hand function through injury or pathology decreases independence, productivity, and quality of life. The scale of this challenge is substantial: hand and wrist injuries are common, occurring at rates of approximately 900 per 100,000 persons annually in the United States and constituting up to 30% of emergency department cases ^1,2^. Neurological conditions compound this burden. Stroke affects an estimated 795,000 Americans annually, ^3^ with upper extremity impairment occurring in up to 88% of cases ^4^, while cerebral palsy affects 1 in 345 children ^5^, and often involves significant hand dysfunction ^6^. The economic and social consequences are severe, with hand injuries alone resulting in millions of days of work absence annually and generating substantial healthcare costs through both direct treatment and productivity losses ^7,8^.

Ultimately, a deeper understanding of how active muscle coordination and passive musculoskeletal properties contribute to dexterous manipulation is needed to effectively advance treatment strategies for hand injury and dysfunction. Computational rigid-body musculoskeletal modeling is a powerful approach and has advanced our understanding of human movement. Musculoskeletal modeling has been extensively applied and validated across multiple domains, as documented in recent reviews of their use in gait analysis ^9^, lower limb biomechanics ^10^, and arm biomechanics^11^. Modeling enables estimation of quantities that are difficult or impossible to measure experimentally, including individual muscle forces, neural excitations of muscle, and joint kinetics^12–16^. These internal variables are fundamental to understanding movement, as they link neural control, muscle coordination, and skeletal mechanics, and ultimately explain how forces are generated and regulated. Beyond advancing fundamental understanding, computational biomechanics approaches have helped improve exoskeleton design ^17^, rehabilitation strategies ^18^, and surgical decision making ^19.^

Unfortunately, comprehensive simulation studies of the full human hand remain scarce for several reasons. Biomechanical models that include all the degrees of freedom (DoF) of the 5 digits of the hand and wrist are particularly rare ^20–23^. Simulating hand motion is computationally demanding due to the hand’s high number of DoFs, small-mass segments that create numerically stiff dynamics, and complex multi-articulate muscle architecture. Furthermore, acquiring detailed validation data is also challenging experimentally due to the hand’s small size and intricate anatomy. As a result, the state of the art for biomechanical simulation of the hand involves a reduced number of fingers or muscles ^24–27^, static optimization ^22,28^, and simplified models of contraction dynamics ^29^.

The objective of this study is to employ state-of-the-art dynamic optimization, direct collocation approaches, which allow us to simulate muscle-driven motions of all five digits of the hand and the wrist. We employ two approaches to evaluate the capabilities and limitations of the simulations: an inverse approach that enforces prescribed kinematics through constraints, and a predictive approach that allows deviations from target trajectories. Additionally, we explore how modifying the finger extensor muscles in a popular open-source musculoskeletal hand model improves simulation accuracy by better reflecting anatomy.

## Materials and Methods

As the basis of our muscle-driven motion simulations of all five digits of the hand and wrist, we collected motion capture data from seven adult participants (4 males, 3 females; age range: 23– 38 years) performing six American Sign Language (ASL) posture matching tasks. These postures were selected to span a wide range of motion across all five fingers. Our human subjects’ study was approved by the Northwestern University Institutional Review Board (Protocols STU00215263 and STU00219081); all participants provided informed consent. Participants were instructed to form a single ASL letter from a resting hand position with the forearm fully supinated. A single trial consisted of the participant repeating this sequence three times in succession for one ASL posture. We collected kinematic data using a custom, 8-camera markerless motion capture system (FLIR BlackFly S GigE RGB cameras with F1.4/6mm and F1.6/4.4-11mm variable focus lenses) sampling at 29 Hz. Joint trajectory reconstruction was performed from video using an end-to-end virtual marker estimation and inverse kinematics pipeline developed by Cotton et. al ^30^ and adapted for hand applications by Firouzabadi et. al^31^. The resulting joint trajectories were segmented into 2-second trials by identifying peaks in joint velocities for task-relevant degrees of freedom. This procedure yielded 255 total trials distributed across six ASL postures (A, B, D, F, L, and O; Fig 1.), with approximately 40–45 trials per posture. Each joint trajectory was then low-pass filtered at 6 Hz with a 3rd order Butterworth filter to remove high-frequency noise. Each trial was time-normalized such that the hand posture at t = 0 *s* corresponded to the rest position, t = 0-1 *s* the transition to the target posture, and t = 1-2 *s* the posture hold phase.

**Figure 1.**
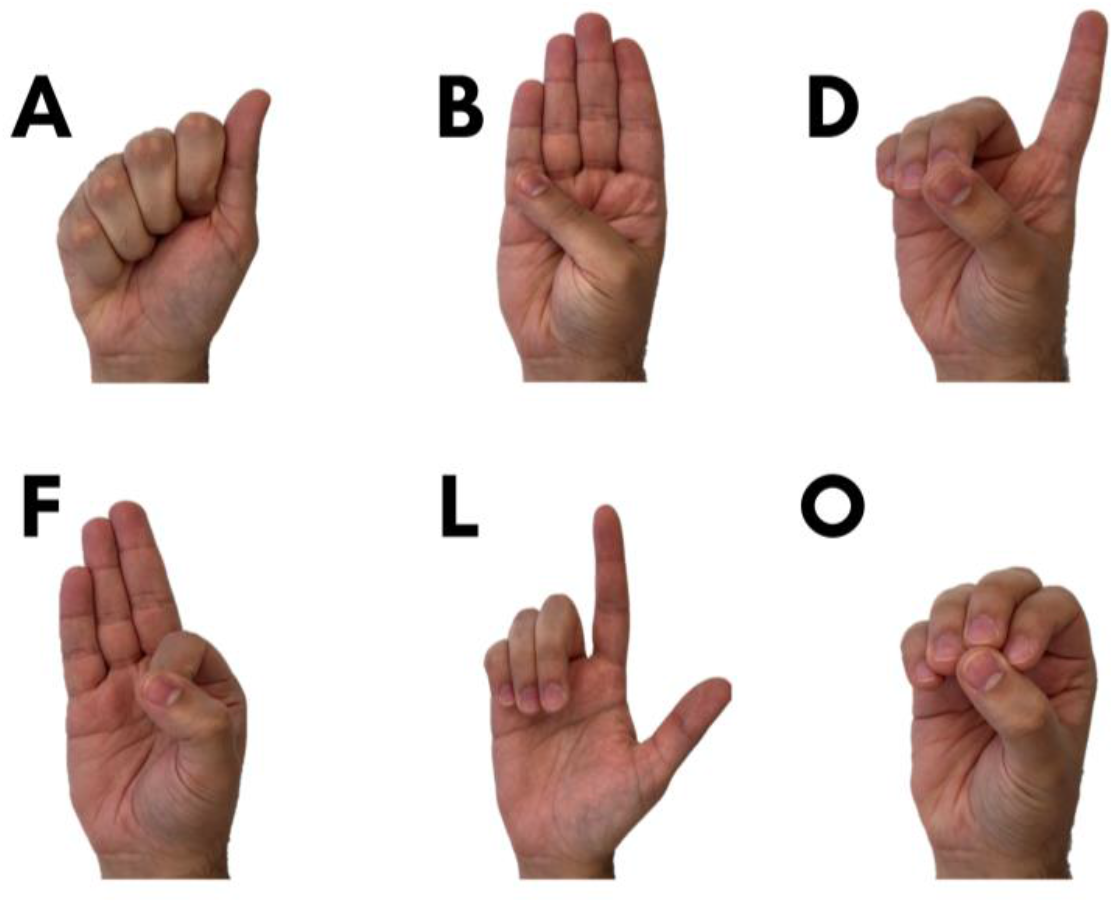
American Sign Language postures used during experimental tasks

Surface EMG recordings of the middle and ring compartments of the superficial finger flexors and finger extensors were collected from one of the seven participants to enable comparison against simulated muscle activations. EMG signals were recorded using a Delsys Bagnoli-16 system (Delsys Incorporated, Natick, MA) at 2048 Hz. Raw EMG signals were band-pass filtered (25-500 Hz), full-wave rectified, low-pass filtered at 10 Hz, and normalized to maximum voluntary contraction (MVC) values obtained during standardized testing procedures.

All dynamic optimization simulations of the hand and wrist motions were performed using OpenSim MOCO^32^. This software provides tools to solve optimal control problems, incorporating the dynamics and constraints of an OpenSim musculoskeletal model. Simulations were performed using the Quest high performance computing facility at Northwestern University. We implemented the biomechanical model of the hand developed by McFarland et. al^22^, which consists of 23 independent DoFs actuated by 43 musculotendon units representing the major intrinsic and extrinsic muscles of the hand and wrist. The wrist is modeled with flexion/extension and radial/ulnar deviation DoFs. Each finger is modeled with flexion/extension at the metacarpophalangeal (MCP), proximal interphalangeal (PIP), and distal interphalangeal (DIP) joints, with abduction/adduction at the MCP joints. The thumb includes carpometacarpal (CMC) flexion/extension and abduction/adduction, MCP flexion/extension, and interphalangeal (IP) flexion/extension. The model also includes a coupled CMC flexion DoF for the fourth and fifth digits, representing the mobility of the ulnar side of the hand. We replaced the muscle model implemented by McFarland et. al^22,33^ with an available muscle model^34^ which provides continuous and differentiable muscle dynamics, essential for gradient-based optimization convergence. To better match the experimentally observed hand orientation, the forearm and elbow posture of the model were set using the rigid body and joint definitions from the upper-limb model of Saul et. al^35^. Although these degrees of freedom were not included as active coordinates in the simulations, the forearm was fixed in maximal supination (−90° pronation–supination in the Saul model convention) and the elbow was fixed at 90° of flexion throughout all simulations.

MOCO uses direct collocation to transform continuous optimal control problems into nonlinear programs by discretizing trajectories into mesh intervals, approximating integrals of cost functions and dynamics via quadrature, and enforcing dynamics as algebraic constraints. This approach naturally incorporates muscle activation dynamics and temporal coupling between time points. Among the collocation schemes available in MOCO, we chose Hermite-Simpson collocation^36^ and solved the resulting nonlinear program using IPOPT ^37^.

MOCO allows target kinematics to be incorporated through two approaches: (1) as constraints that enforce tracking within a specified tolerance (MocoInverse), or (2) integrated into the cost function as tracking terms that minimize deviations from reference trajectories (MocoTrack). Here we used both approaches (as outlined in Fig. 2). We initially used MocoInverse due to its computational efficiency. Our results from these simulations motivated a subsequent set of simulations using the MocoTrack formulation. Solver convergence statistics for both approaches are reported in Supplementary Table 1.

**Figure 2.**
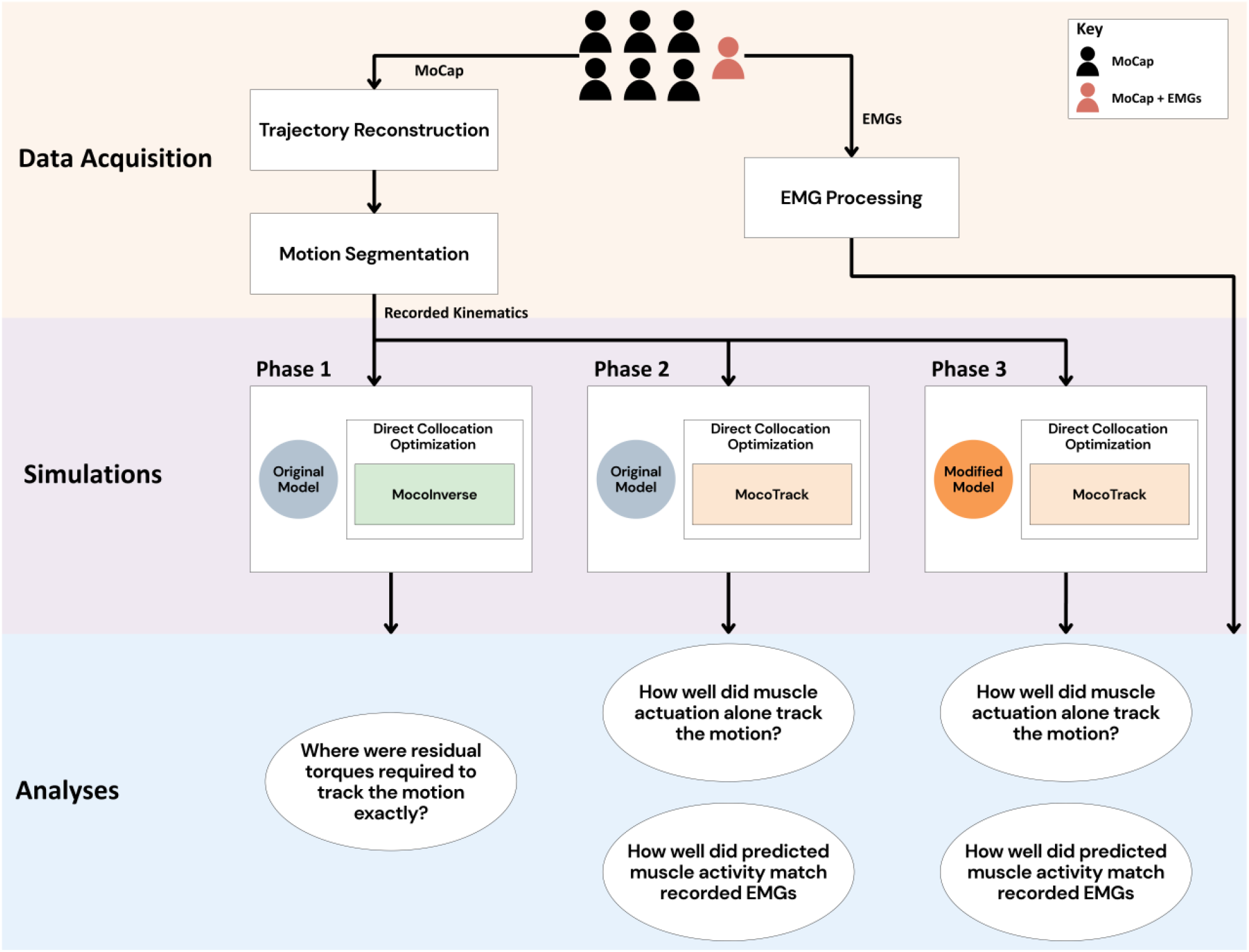
Multi-phase computational pipeline for evaluating muscle-driven motion tracking. Motion capture (MoCap) data were collected from seven participants performing posture matching tasks, with one participant additionally instrumented with EMG sensors. Following trajectory reconstruction and motion segmentation, recorded kinematics served as inputs to three sequential simulation phases. Phase 1 employed direct collocation with MocoInverse using the original musculoskeletal model to determine residual torques required for exact motion reproduction. Phase 2 used MocoTrack with the same model to assess how well muscle actuation alone could track the observed motions and predict muscle activity patterns. Phase 3 applied MocoTrack to a modified musculoskeletal model to evaluate whether targeted model refinements improved both motion tracking accuracy and EMG prediction fidelity.

### Inverse Formulation (MocoInverse)

Our initial simulation approach implemented MocoInverse to solve for the muscle activations required to replicate the recorded hand and wrist kinematic trajectories within a strict tolerance (1×10^−3^ degrees, the default tolerance setting in MocoInverse). This constraint-based approach prescribed the observed motion, eliminating the need to integrate multibody dynamics and substantially reducing computational cost. However, because of the strict requirements imposed by these kinematic constraints, idealized torque actuators (commonly termed “reserve actuators”) were included to supplement muscle forces and improve convergence feasibility. Such reserve actuators are standard in the OpenSim simulation tool-suite to actuate joints directly or to handle dynamic inconsistencies between experimental data and models, which can arise from measurement noise, modeling simplifications, or insufficient actuator capacity^38,39^. For our MocoInverse simulations, reserve actuators were included at each joint in the hand and wrist. The optimization problem was formulated to minimize a cost function comprising the sum of squared muscle activations and reserve actuator controls (i.e., an effort-minimization term). To discourage reliance on reserve actuators, the reserve control terms were weighted 1000 times higher in the cost function.

OpenSim’s Muscle Analysis tool was used to compute muscle-induced joint moments, defined as the rotational torques generated by each muscle about a joint given its activation and anatomical configuration. Muscle contribution to joint torque was quantified as the percentage of the total joint torque attributable to muscle-induced joint moments, relative to the combined contributions of muscles and reserve actuators. Specifically:

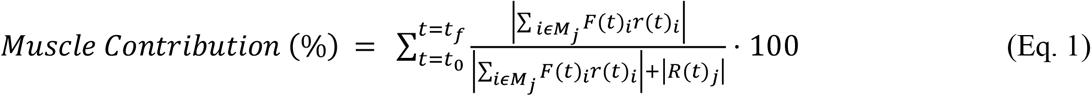

Where:

*M*_*j*_ **-** Set of muscles that generate torques about joint *j*

*F*_*i*_ **-** Force generated by muscle *i*

*r*_*i*_ - Moment arm of muscle *i* with respect to joint *j*

*R*_*j*_ - Reserve Actuator Torque at joint *j*

*t*_0_ - Initial time of trajectory

*t*_*f*_ - Final time of trajectory

Higher values of this metric indicate greater reliance on muscle actuation, and lower values reflect greater dependence on artificial torque supplementation. Torque contributions from the artificial torque actuators were calculated directly from their solved-for controls.

### Predictive Formulation (MocoTrack)

To evaluate whether the hand model could reproduce experimentally measured kinematics using only physiological muscle forces, without supplementation from reserve actuators, we implemented MocoTrack. MocoTrack is considered a predictive simulation approach because it jointly solves for muscle activations and kinematics simultaneously, rather than prescribing the motion a priori. Unlike MocoInverse, which enforces experimental kinematics as strict constraints, MocoTrack incorporates kinematic tracking errors into the cost function, allowing the model to deviate from experimental kinematics when necessary to satisfy dynamic feasibility with muscle-only actuation. However, this added flexibility comes at a computational cost, as multibody dynamics are now part of the optimization problem, requiring integration of the equations of motion to solve for kinematic trajectories alongside muscle control.

We tested multiple cost function formulations to examine how tracking priorities influenced tracking error and predicted muscle activity. Simulations were first performed using (1) kinematic tracking terms alone and (2) kinematic tracking combined with a sum-of-squared muscle activation penalty (applied only to the single subject with available EMG data). Then, we selectively increased tracking error weights for the wrist, DIP, or PIP joints to assess how prioritizing tracking at specific joints affected global tracking behavior and muscle activation patterns.

Simulation performance was evaluated by computing the mean absolute joint angle error for each joint relative to the experimental kinematic trajectory. We compared mean absolute tracking errors across joints using non-parametric statistics due to non-normal distributions. (Shapiro-Wilk, p < 0.0001) and heterogeneous variances (Levene’s test, p < 0.0001) in the tracking error data. Effect sizes were quantified using rank-biserial correlation (|r| < 0.1: negligible; 0.1-0.3: small; 0.3-0.5: medium; ≥0.5: large).

Based on observations from MocoInverse simulations, we hypothesized that MocoTrack tracking errors differed across joint types (MCP, PIP, DIP) within individual fingers. For each finger, we applied Kruskal-Wallis H-tests to assess overall differences among joints (H_0_: tracking errors equivalent across all joints; H_1_: at least one joint differs), followed by pairwise Mann-Whitney U tests to compare tracking errors between each pair of joints, with false discovery rate correction via the Benjamini-Hochberg procedure (α = 0.05).

### Anatomical Refinement and Model Validation

#### Dual-Insertion EDC Model Development

To address limitations observed in the simulation results observed with the original model, we developed a modified version incorporating an anatomically refined representation of the extensor digitorum communis (EDC) muscle to better capture the complexity of the finger extensor mechanism. Hereafter, we refer to this modified model as the dual-insertion EDC model. In addition to the original EDC compartments (EDCI, EDCM, EDCR, and EDCL, where the suffixes I, M, R, and L denote the index, middle, ring, and little finger compartments, respectively) that insert at the distal phalanges, we added four central slip musculotendon units (EDCI_CS, EDCM_CS, EDCR_CS, and EDCL_CS) that insert at the middle phalanges to capture the EDC’s dual insertion pattern. Since these represent anatomical splits of single tendons rather than independent muscles, we implemented OpenSim SynergyControllers to constrain each EDC compartment and its corresponding central slip to share identical activation patterns (e.g., EDCM and EDCM_CS). The maximum isometric force capacity was redistributed between the distal and central slip insertions based on physiological data, with 63% allocated to the original distal insertion units and 37% to the newly added central slip units, reflecting the relative force-generating capacity of each insertion site^40^.

To compare the impacts of these modifications to the original model, we ran the MocoTrack simulations with and without this modified model. We performed paired Wilcoxon signed-rank tests to test whether the dual-insertion EDC model reduced mean absolute joint tracking errors compared to the original model. To control for multiple comparisons across all tested joints, we applied the Benjamini-Hochberg false discovery rate correction to the p-values, with adjusted p-values < 0.05 considered to be statistically significant.

## Results

### MocoInverse Simulations

#### A. Muscle Contribution Analysis

When constrained to reproduce the experimental kinematic trajectories within a strict tolerance (using MocoInverse), the wrist achieved nearly complete muscle-driven control, with muscle forces contributing 96-99% of the control of radiocarpal flexion and deviation respectively (Fig. 3). In contrast, many finger and thumb joints required supplementary torques to bridge the gap between simulated muscle-driven motion and the prescribed kinematics (Fig. 3). The thumb, index, and middle fingers showed a proximal-to-distal gradient in the contribution of supplementary torques, with more distal joints requiring greater supplementary torques to achieve the desired motions. Results were highly consistent among all PIP and DIP joints, where contributions from reserve torque actuators were approximately equal to those from muscles, regardless of digit. Except for the coupled 4^th^ and 5^th^ carpometacarpal (CMC) flexion DoF^41^, muscle contributions were nearly equal to or greater than those from supplementary torque actuators. The average patterns across all trials were consistent among all six ASL postures (Supplementary Figure S1).

**Figure 3.**
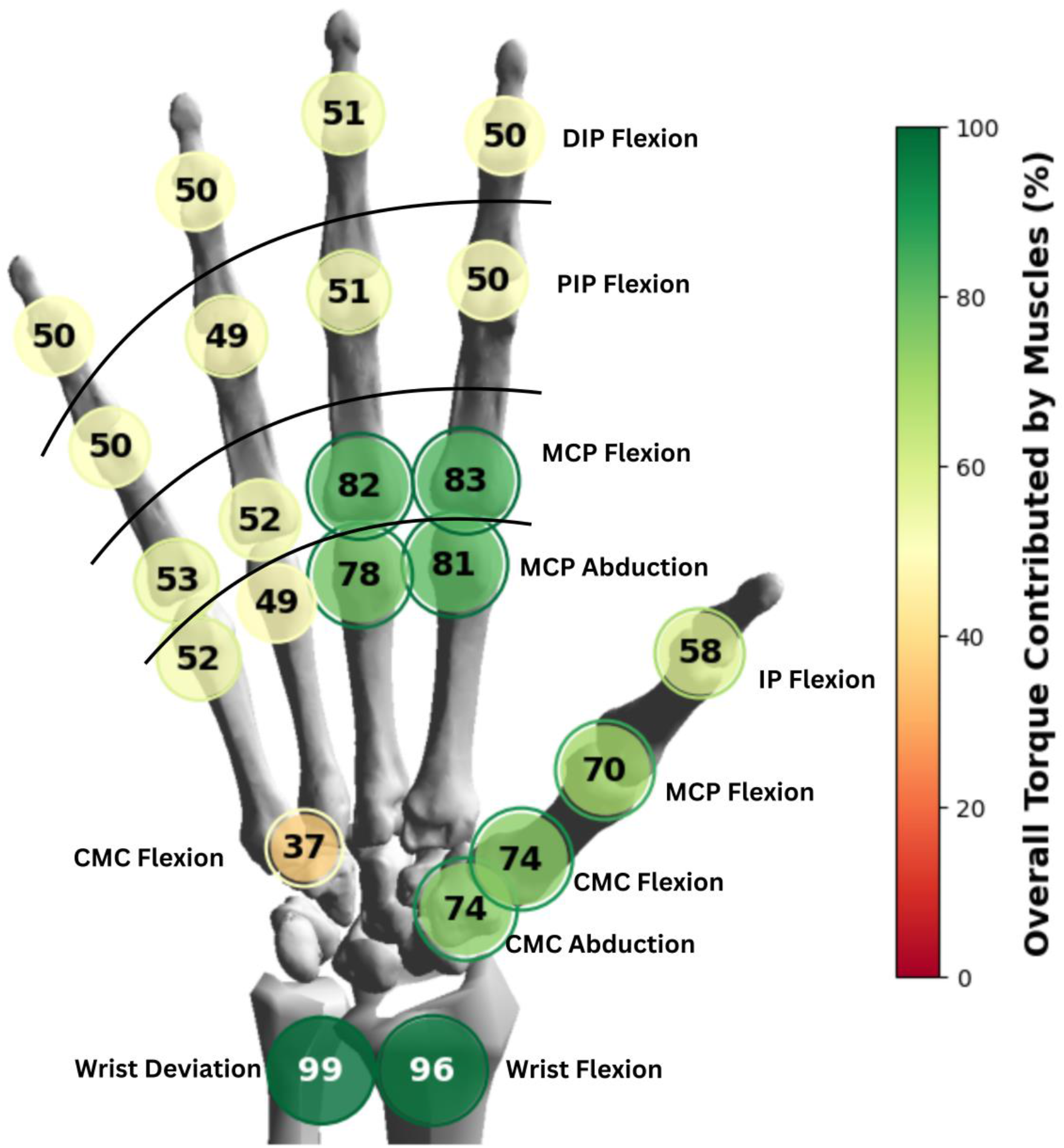
Visual representation of muscle versus reserve actuator contributions to total joint torque at each degree of freedom in the hand and wrist, computed using MocoInverse. Each degree of freedom is marked with a colored circle that indicates the relative muscle contribution at that joint. The full color gradient spans dark red (none to very limited muscle contribution to total joint torque) to dark green (primarily muscles contributed to total joint torque); results are averaged across time, trials, postures, and participants. A separate ring, with diameter equal to the mean muscle contribution + 1 standard deviation (s.d.) surrounds each filled circle. The color of the s.d. ring also indicates the value of the mean + 1 s.d. for that degree of freedom. In general, muscle contributions decreased from proximal-to-distal.

### MocoTrack Predictive Simulations

#### B. Tracking Performance

When we used the predictive simulation approach (MocoTrack) to solve for muscle activations without prescribing motion a priori and prioritized wrist tracking, DIP and PIP flexion exhibited the largest tracking errors across all four fingers (p < 0.001; Fig. 4). Notably, experimental motion amplitude differed substantially across joints: peak-to-peak excursion was modest at the wrist and MCP abduction (∼3–12°), but much greater for MCP flexion (∼33–37°) and for PIP and DIP flexion (up to ∼55°). In the baseline tracking formulation (equally weighted kinematic tracking terms for all joints), wrist flexion and deviation tracking errors were large relative to their modest excursions (wrist flexion MAE ≈ 16.14°; wrist deviation MAE ≈ 13.83°). Increasing the wrist tracking weight in the cost function by 10 reduced these errors by approximately 90% (from 16.14° to 1.64° for wrist flexion and from 13.83° to 1.74° for deviation), while maintaining the tracking error at other joints to within ∼2°.

**Figure 4.**
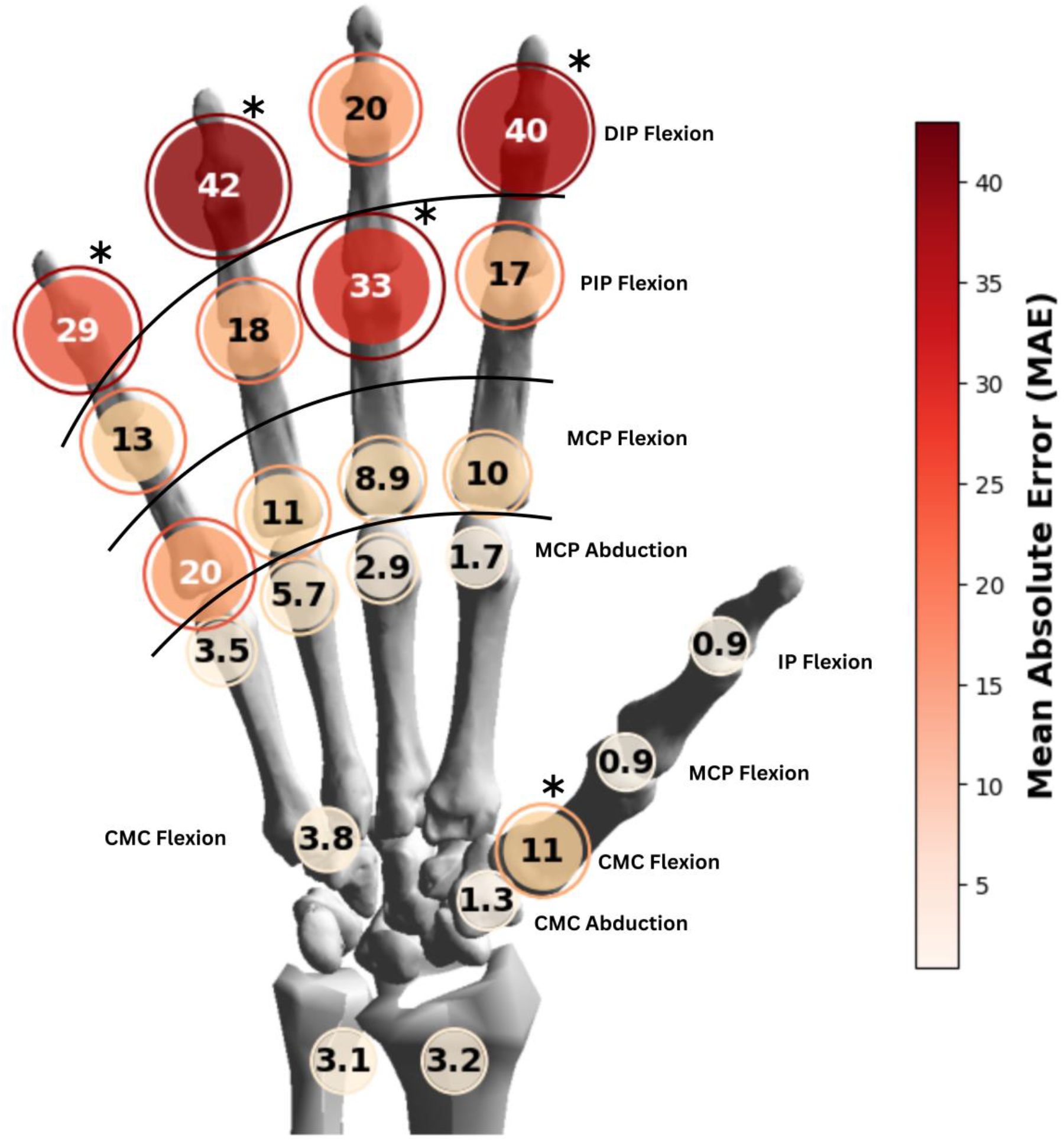
Kinematic tracking errors, computed as the mean absolute joint angle error averaged across time, trials, postures, and participants from MocoTrack simulations. Mean error magnitude (in degrees) is reported directly by the text within each filled circle and is symbolically displayed by both circle size and color intensity; larger, darker circles indicate greater tracking errors. A separate ring, with diameter and color equal to the value of the mean error + 1 s.d. surrounds each filled circle. For each digit, the joint with the largest error (p < 0.001) is labeled; an asterisk indicates the single joint that had the largest error for that digit.

Simulated DIP finger joint trajectories were consistently more flexed than experimental measurements. For example, for task L, DIP flexion angles for the ring, middle, and index fingers were offset by 36.1±13.6° in the flexion direction throughout the entire movement (Fig. 5). This flexion bias was consistent across tasks, as reflected by the outward-pointing spikes at the DIP joints in the task-averaged tracking error (Fig. 6) and in the task-specific profiles (Supplementary Fig. S2).

**Figure 5.**
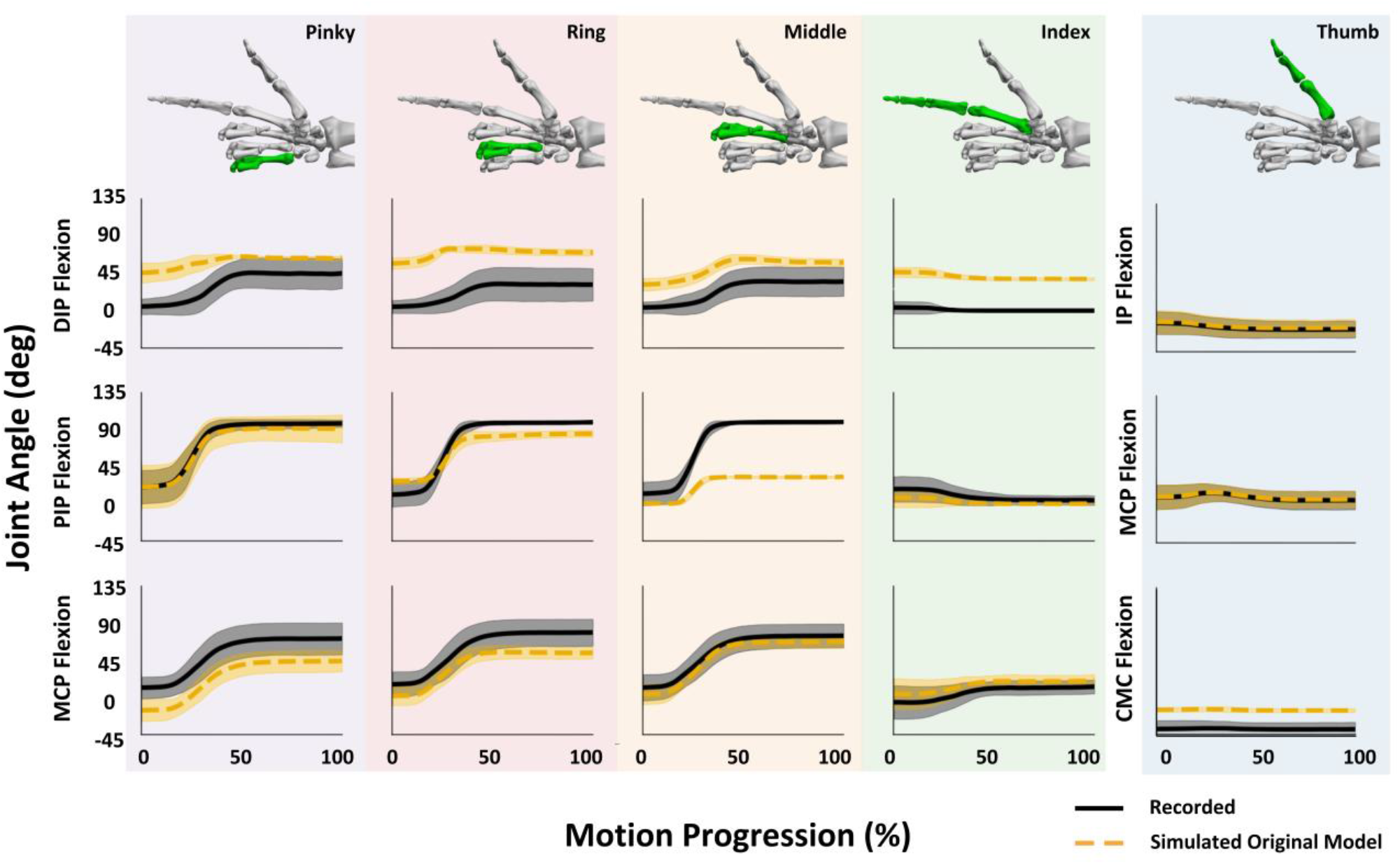
Comparison of experimental (solid black lines) and simulated (dashed orange lines) joint kinematics for the ASL letter L task. Joint angle trajectories are shown across time-normalized movement progression (0–100%). Each column represents a digit (Pinky through Thumb, left to right), color-coded by digit for visual reference, with rows representing joint degrees of freedom. Shaded regions indicate ±1 standard deviation across converged trials for experimental (gray) and simulated (orange) data. Positive angles indicate flexion. All trajectories are averaged across 28 converged trials.

**Figure 6.**
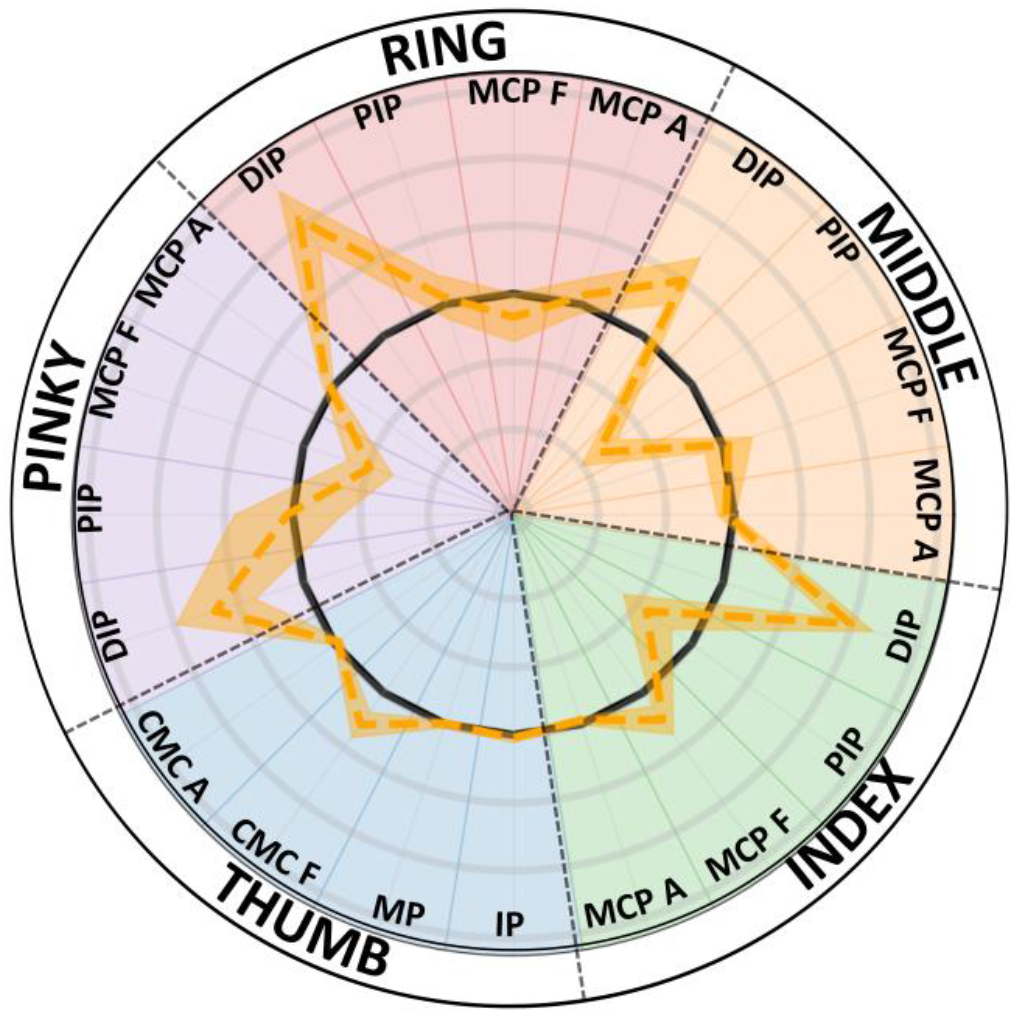
Tracking error across hand and thumb joints averaged across all ASL tasks. The orange dashed line represents the signed mean tracking error for each degree of freedom, averaged across all converged trials and participants; shaded regions indicate ±1 standard deviation. The solid black line denotes perfect tracking (0° error). Positive values (outward from 0°) indicate errors in the flexion or abduction direction depending on the joint, whereas negative values (inward) indicate extension or adduction errors. Background colors group joints by digit and match the color scheme used in Figure 5. Light gray concentric rings indicate 20° radial increments. A consistent outward spike is observed at the DIP joint of each finger, reflecting a flexion bias across tasks, whereas other joints exhibit smaller errors with more variable directional patterns.

Unexpectedly, the model exhibited a strong bias toward extensor activation with minimal flexor engagement across tasks. This pattern persisted even when the simulations minimized overall control effort. Across all six ASL tasks examined (A, B, D, F, L, O), the model consistently predicted high mean EDCM activity (41–55% of maximal activation, averaged across trials and timepoints), whereas activation of the middle finger flexor digitorum superficialis (FDS) remained negligible (<0.01%). For example, during the flexion-dominant ASL A task, the model predicted substantial EDCM activity (55±13% maximal activation) with minimal FDS activity (∼0.004%), whereas EMG recordings showed low extensor activity (3±1% MVC) and flexor activity that, although modest, exceeded extensor activation (8±4% MVC). In the extension-dominant ASL B task, predicted EDCM activity remained similarly elevated (54±10% maximal activation) with negligible FDS activity (∼0.006%), whereas EMG recordings again showed substantially lower extensor activity (13±6% MVC) and small but measurable flexor activity (2±1% MVC) (Fig. 7).

**Figure 7.**
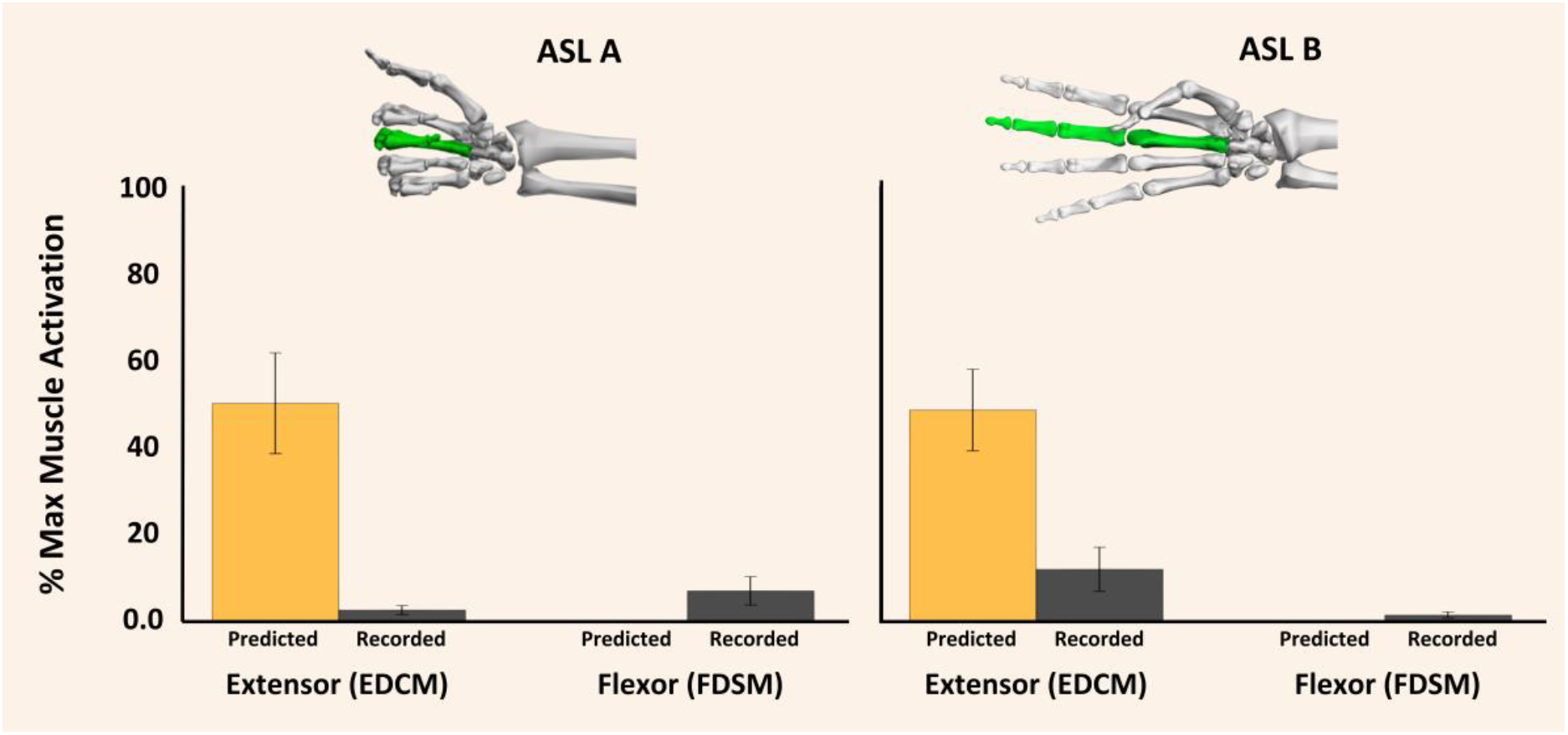
Comparison of predicted and recorded muscle activation levels for middle finger extensor and flexor muscles during ASL posture matching tasks. Mean muscle activation levels (±1 s.d.) expressed as percentage of max activation are shown for the middle compartment of the extensor digitorum communis (EDCM) and the flexor digitorum superficialis (FDSM) muscles across all trials and timepoints for one subject with available surface EMG data. Two American Sign Language (ASL) letter tasks are shown: ASL A (left panel), which requires primarily middle finger flexion, and ASL B (right panel), which requires primarily middle finger extension. Orange bars represent MocoTrack predicted mean muscle activations as percentage of maximal activation. Black bars represent recorded EMG signals as a percentage of maximum voluntary contraction (MVC). Following the digit color-coding scheme from previous figures, the tan background indicates the middle finger column, and hand skeletal models above each panel show this same finger highlighted in green with the hand posed in the final target posture for each respective ASL letter.

### Dual Insertion Extensor Model Simulations

Overall, using the dual-insertion EDC model led to widespread reductions in tracking error across the hand, with the most consistent improvements observed at the DIP joints (Fig. 8). DIP tracking errors decreased significantly across all digits (False Discovery Rate-adjusted p < 0.001), with reductions ranging from approximately 10–65% and averaging ∼35% overall. PIP flexion errors also decreased significantly for the index and middle fingers. MCP flexion errors were within ∼3° of the original simulations. MCP abduction degrees of freedom exhibited small but statistically significant reductions in tracking error (generally <3°).

**Figure 8.**
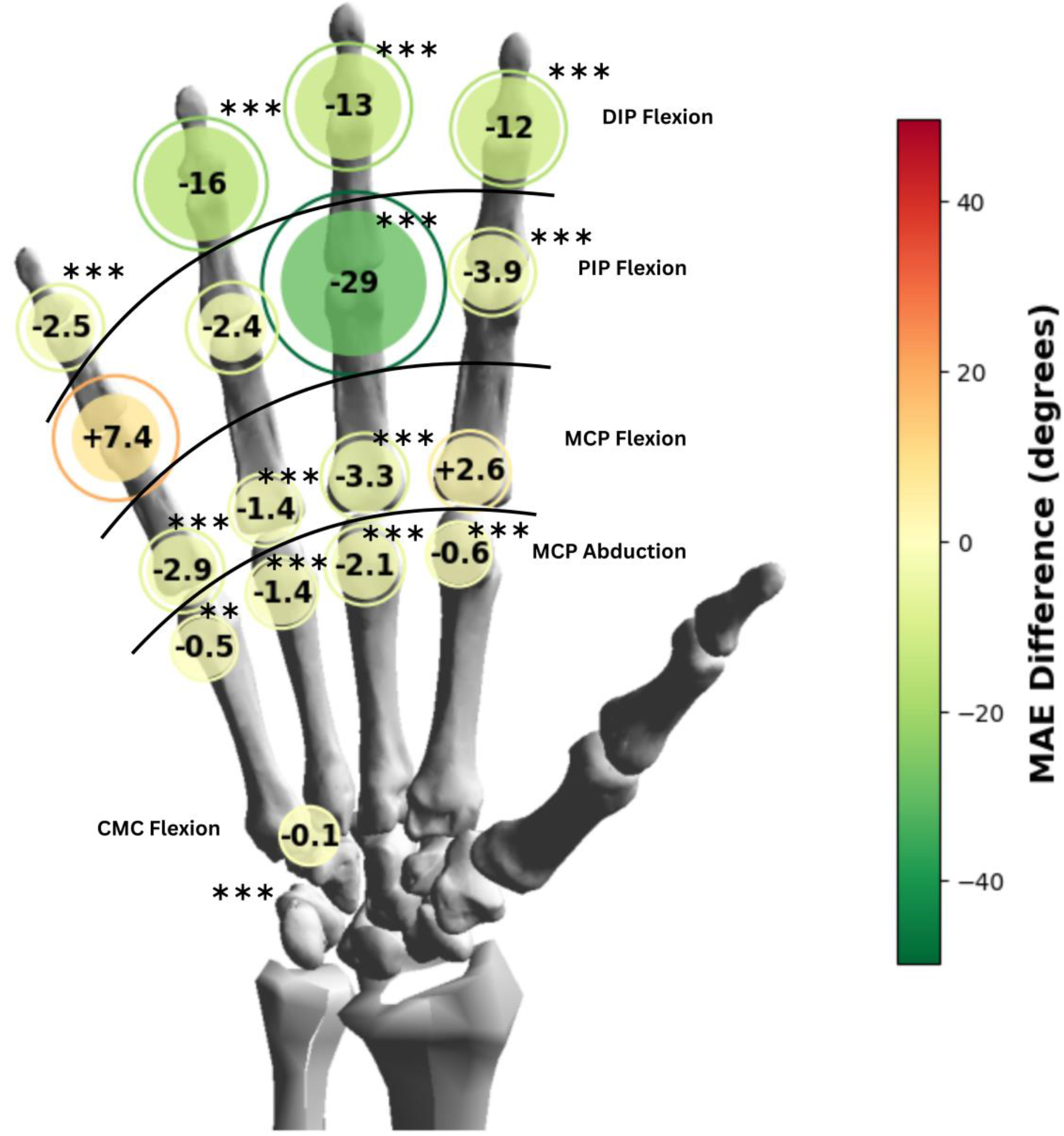
Mean differences in joint angle tracking errors between the dual-insertion and original EDC models, computed from MocoTrack simulations and averaged across time, trials, and postures. The difference (in degrees), including sign, is reported directly by the text within each filled circle and is symbolically displayed by both circle size and color on a diverging red-yellow-green gradient centered at zero, where circle size reflects difference magnitude and color reflects both sign and magnitude. Negative values (green) indicate that the dual-insertion model had lower tracking errors than the original model. Positive values (red) indicate higher errors. For negative differences, an outer ring with diameter corresponding to the mean difference minus 1 standard deviation is displayed. For positive differences, an outer ring with diameter corresponding to the mean difference plus 1 standard deviation is displayed. The color of each ring is consistent with the diverging color gradient at its respective value. Statistically significant decreases in error are indicated by asterisks: ** p < 0.01, *** p < 0.001.

### Muscle Activation Patterns

The modified EDC model reduced predicted extensor activation levels for the middle finger compartment, the only digit with available EMG data for validation. For the middle finger during the flexion-dominant ASL A task, the modified model reduced predicted EDCM activation by approximately 75% (from 55±13% to 14±9% maximal activation), compared to recorded EMG levels of 3±1% MVC. For the extension-dominant ASL B task, predicted EDCM activation decreased by approximately 38% (from 54±10% to 33±4% maximal activation), compared to EMG recordings of 13±6% MVC (Fig. 9). While the modified model substantially reduced predicted EDCM activation relative to the original model, discrepancies remained: predicted activation exceeded EMG by approximately 4-fold for ASL A and 2.5-fold for ASL B. FDSM activation remained negligible in both models relative to recorded EMG. Although the modified model increased predicted FDSM activation slightly during ASL A (from ∼0.004% to ∼0.17% maximal activation), values remained orders of magnitude lower than measured EMG levels (8±4% MVC). Similar discrepancies were observed during ASL B, where predicted FDSM activation remained near zero despite measurable flexor EMG activity (2±1% MVC).

**Figure 9.**
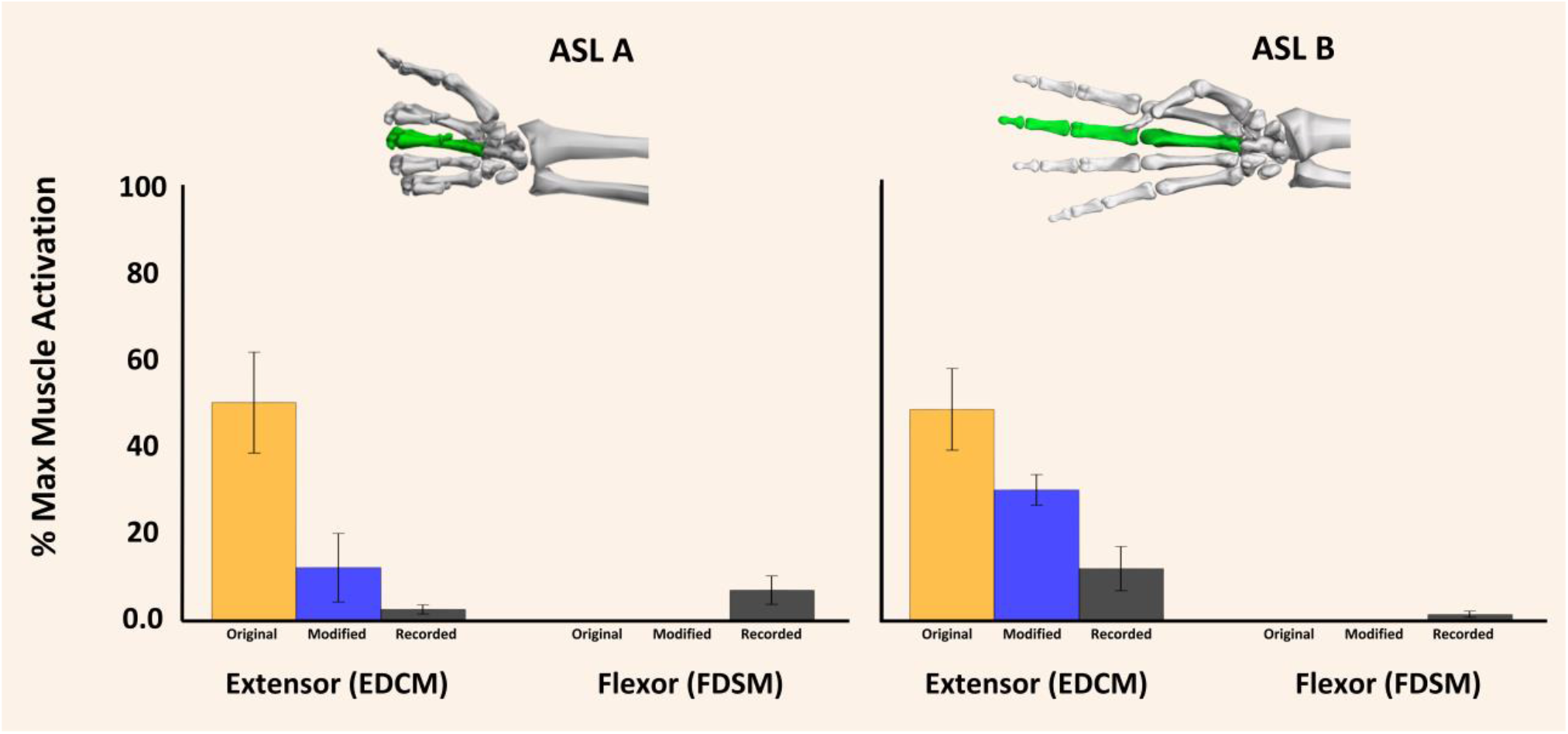
Comparison of original model, modified model, and recorded muscle activation levels for middle finger muscles during ASL tasks. Mean muscle activation levels (±SD) expressed as percentage of max activation are shown for extensor digitorum communis middle (EDCM) and flexor digitorum superficialis middle (FDSM) across all trials and timepoints for one subject with available surface electromyography (EMG) data. Two American Sign Language (ASL) letter tasks are shown: ASL A (left panel, flexion-dominant) and ASL B (right panel, extension-dominant). Orange bars represent the original model predictions as percentage of maximal activation, blue bars represent the modified dual-insertion EDC model predictions as percentage of maximal activation, and black bars represent recorded EMG signals as percentage of maximum voluntary contraction (MVC). Following the digit color-coding scheme from previous figures, the tan background indicates the middle finger column, and hand skeletal models above each panel show this same finger highlighted in green with the hand posed in the final target posture for each respective ASL letter.

## Discussion

This study demonstrated the application of direct collocation methods to full-hand simulations of multi-joint postures. Different optimization formulations provided complementary diagnostic information that helped identify specific anatomical limitations in the model. MocoInverse simulations showed a proximal-to-distal gradient in the supplementary torque required to bridge the gap between simulated muscle-driven motion and the prescribed kinematics. The reliance on reserve actuators in MocoInverse simulations, particularly at distal joints, suggested that the current model’s muscle architecture could not generate the precise force combinations required to replicate experimental kinematics. Notably, reserve actuators were often observed compensating for muscle-generated joint moments that exceeded those required to reproduce the target kinematics at individual joints, indicating that the underlying issue was not insufficient force generation but rather limited coordination of muscle forces across multiple joints. When we removed the prescribed kinematics and solved the problem using the predictive MocoTrack formulation, the simulations produced similar patterns, further indicating coordination limitations at the distal joints. Consistent with this interpretation, when we selectively increased tracking priorities for the distal joints, we observed a trade-off between PIP and DIP tracking, with improvements at one joint typically accompanied by increased errors at the other.

Based on these findings, we hypothesized that known simplifications in extensor representation underlie the coordination limitations observed in our simulations. The extensor hood is a complex passive structure that coordinates extension forces across the PIP and DIP joints. The EDC tendon splits into a central slip, which inserts at the middle phalanx to extend the PIP joint, and two lateral bands that converge and insert at the distal phalanx to extend the DIP joint. Through these lateral bands, the extensor hood also integrates intrinsic muscles (lumbricals and interossei) with the EDC, creating a distributed force transmission system that enables coordinated extension across the finger joints^42^. However, current rigid body full hand models employ simplified representations of the EDC that do not fully capture these interactions or the distribution of extensor forces across the finger joints. To test this hypothesis, we developed a dual-insertion EDC model that explicitly represented the central slip insertion at the middle phalanx alongside the existing distal insertion. This modification significantly improved kinematic tracking performance, particularly at the DIP and PIP joints where the original model showed the largest deficiencies. The architectural refinement also reduced predicted activation magnitudes in the middle and ring finger EDC compartments toward experimentally measured EMG values, although discrepancies between predicted and recorded activations remained.

### Limitations and Future Directions

Several limitations of this study warrant consideration. First, our experimental protocol focused on posture matching tasks only. While these postures span a wide range of joint configurations, future work should examine whether the dual-insertion EDC model improves predictions during functional tasks involving object interaction and manipulation. Second, EMG validation was limited to a single participant and focused only on the superficial finger extensors and flexors. Expanding EMG recordings to include more muscles and additional participants would provide more comprehensive validation of predicted muscle activation patterns. Third, the dual-insertion EDC model represents only one step toward a more complete representation of the extensor hood mechanism. More detailed biomechanical models of the finger extensor apparatus have been proposed that explicitly represent lateral bands, intrinsic muscle insertions, and the interconnected tendon network^43–45^. These approaches capture the distributed force transmission within the extensor mechanism more faithfully than the simplified tendon representations commonly used in multibody musculoskeletal models. However, incorporating these structures into whole-hand optimal control simulations remains challenging because the extensor hood functions as a branching tendon network rather than a set of independent musculotendon paths. Representing these interactions typically requires additional constraints or continuum mechanics formulations, which substantially increase model complexity and computational cost.

## Conclusion

Direct collocation methods enable muscle-driven simulations of full-hand movements but reveal critical limitations in current hand model representations. The systematic tracking errors and reserve actuator reliance observed across complementary optimization formulations pointed specifically to inadequate representation of the extensor mechanism. A dual-insertion EDC model that better captures the anatomy of the extensor hood significantly improved kinematic tracking, while bringing predicted muscle activations closer to experimental EMG recordings. These findings demonstrate that direct collocation methods, when combined with anatomically informed model refinements, show promise for achieving physiologically realistic muscle-driven hand simulations. Further development of comprehensive hand models incorporating detailed representations of passive structures and intrinsic-extrinsic muscle interactions will be essential for translational applications in surgical planning, rehabilitation, and assistive device design.

## Acknowledgements

Research reported in this publication was supported by the National Institute Of Neurological Disorders And Stroke of the National Institutes of Health under Award Number R01NS131953. The content is solely the responsibility of the authors and does not necessarily represent the official views of the National Institutes of Health. Research reported in this publication was supported by the Eunice Kennedy Shriver National Institute Of Child Health & Human Development of the National Institutes of Health under Award Number T32HD007418. This research was supported in part through the computational resources and staff contributions provided for the Quest high performance computing facility at Northwestern University which is jointly supported by the Office of the Provost, the Office for Research, and Northwestern University Information Technology.

## AI Use Statement

The authors used ChatGPT (OpenAI, GPT-5.3) to assist with language editing and drafting revisions. All outputs were reviewed and verified by the authors, who take full responsibility for the final manuscript.

## Disclosure Statement

The authors report there are no competing interests to declare.

## Author Contributions Statement

M.C. conceived and designed the study, developed the computational model, performed simulations and analysis, and drafted the manuscript. P.F., F.R., and V.D. contributed to experimental data collection. P.F. and R.J.C. developed motion capture tools used in the study, and F.R. contributed software for EMG data processing. R.J.C. provided methodological support and resources for motion capture data collection. L.E.M. and W.M.M. supervised the project and critically revised the manuscript. W.M.M. also contributed to study conceptualization and project oversight. All authors reviewed and approved the final manuscript and agreed to be accountable for all aspects of the work.

## Supplementary Material

**Table S1.** Convergence rates across simulations for each experimental posture. Entries report on the number of successful optimizations relative to the total attempted (percentage in parentheses). For comparisons between the two models, only the same converged trials were used.

| Model | Cost | Method | A | B | D | F | L | O | Overall |
| --- | --- | --- | --- | --- | --- | --- | --- | --- | --- |
| Original | Effort Penalty | MocoInverse | 19/42<br>(45%) | 22/41<br>(54%) | 28/45<br>(62%) | 24/42<br>(57%) | 26/44<br>(59%) | 20/41<br>(49%) | 139/255<br>(55%) |
| Original | Balanced Tracking | MocoTrack | 36/42<br>(86%) | 40/41<br>(98%) | 44/45<br>(98%) | 42/42<br>(100%) | 36/44<br>(82%) | 39/41<br>(95%) | 237/255<br>(93%) |
| Original | Strong Wrist Tracking | MocoTrack | 36/42<br>(86%) | 40/41<br>(98%) | 44/45<br>(98%) | 42/42<br>(100%) | 36/44<br>(82%) | 39/41<br>(95%) | 237/255<br>(93%) |
| Original | Strong DIP Tracking | MocoTrack | 37/42<br>(88%) | 31/41<br>(76%) | 42/45<br>(93%) | 30/42<br>(71%) | 35/44<br>(80%) | 33/41<br>(80%) | 208/255<br>(82%) |
| Original | Strong PIP Tracking | MocoTrack | 27/42<br>(64%) | 35/41<br>(85%) | 38/45<br>(84%) | 42/42<br>(100%) | 36/44<br>(82%) | 37/41<br>(90%) | 215/255<br>(84%) |
| Dual-Insertion EDC | Balanced Tracking | MocoTrack | 40/42<br>(95%) | 40/41<br>(98%) | 44/45<br>(98%) | 41/42<br>(98%) | 42/44<br>(95%) | 39/41<br>(95%) | 246/255<br>(96%) |
| *Original | Strong Wrist Tracking + Effort Penalty | MocoTrack | 5/6<br>(83%) | 6/6<br>(100%) | 6/6<br>(100%) | 6/6<br>(100%) | 6/6<br>(100%) | 6/6<br>(100%) | 35/36<br>(97%) |

\*Simulations were run only for the subject with available EMG data.

**Figure S1.**
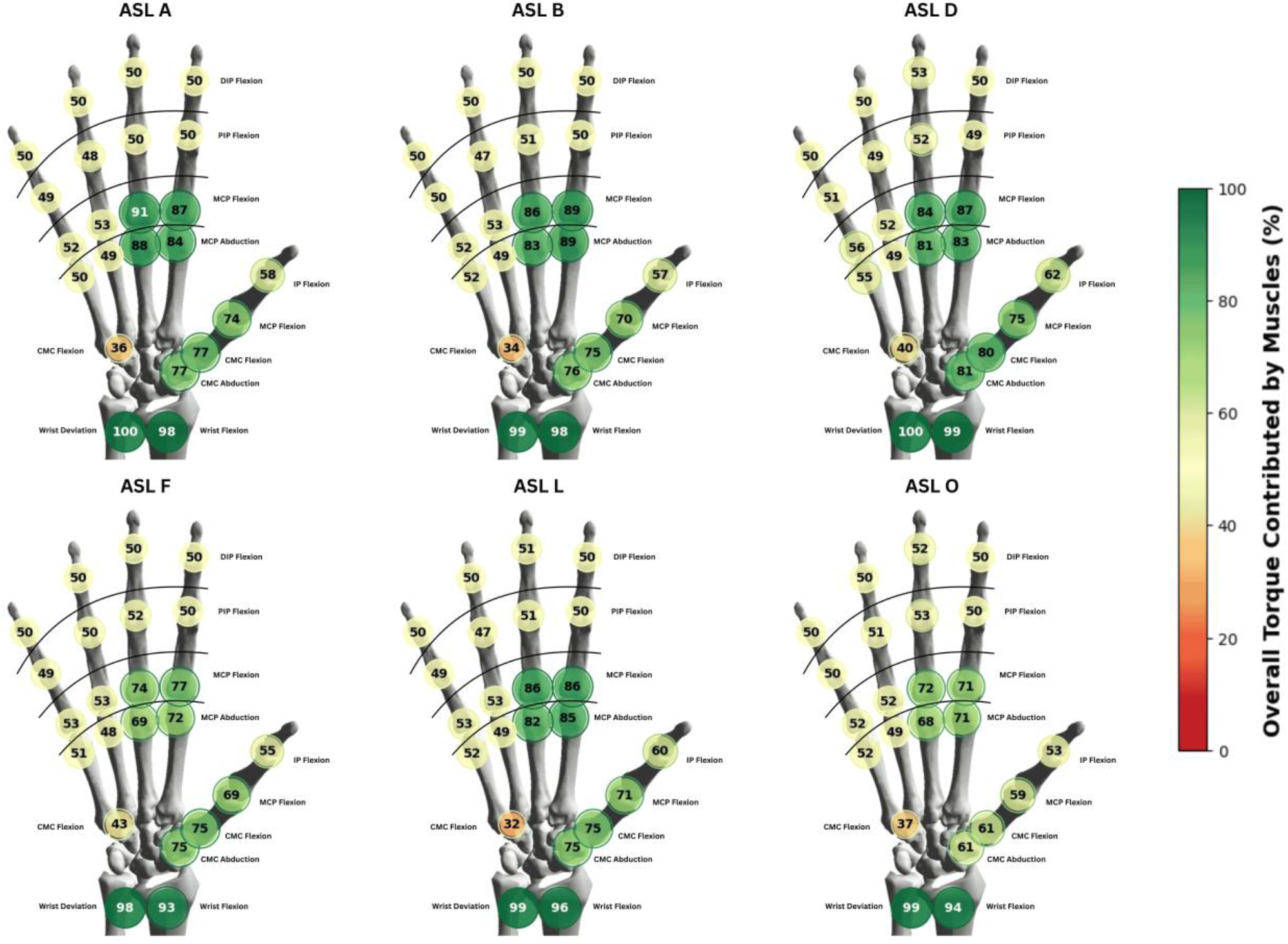
Visual representation of muscle versus reserve actuator contributions to total joint torque at each degree of freedom in the hand and wrist, computed using MocoInverse for all 6 ASL posture matching tasks.

**Figure S2.**
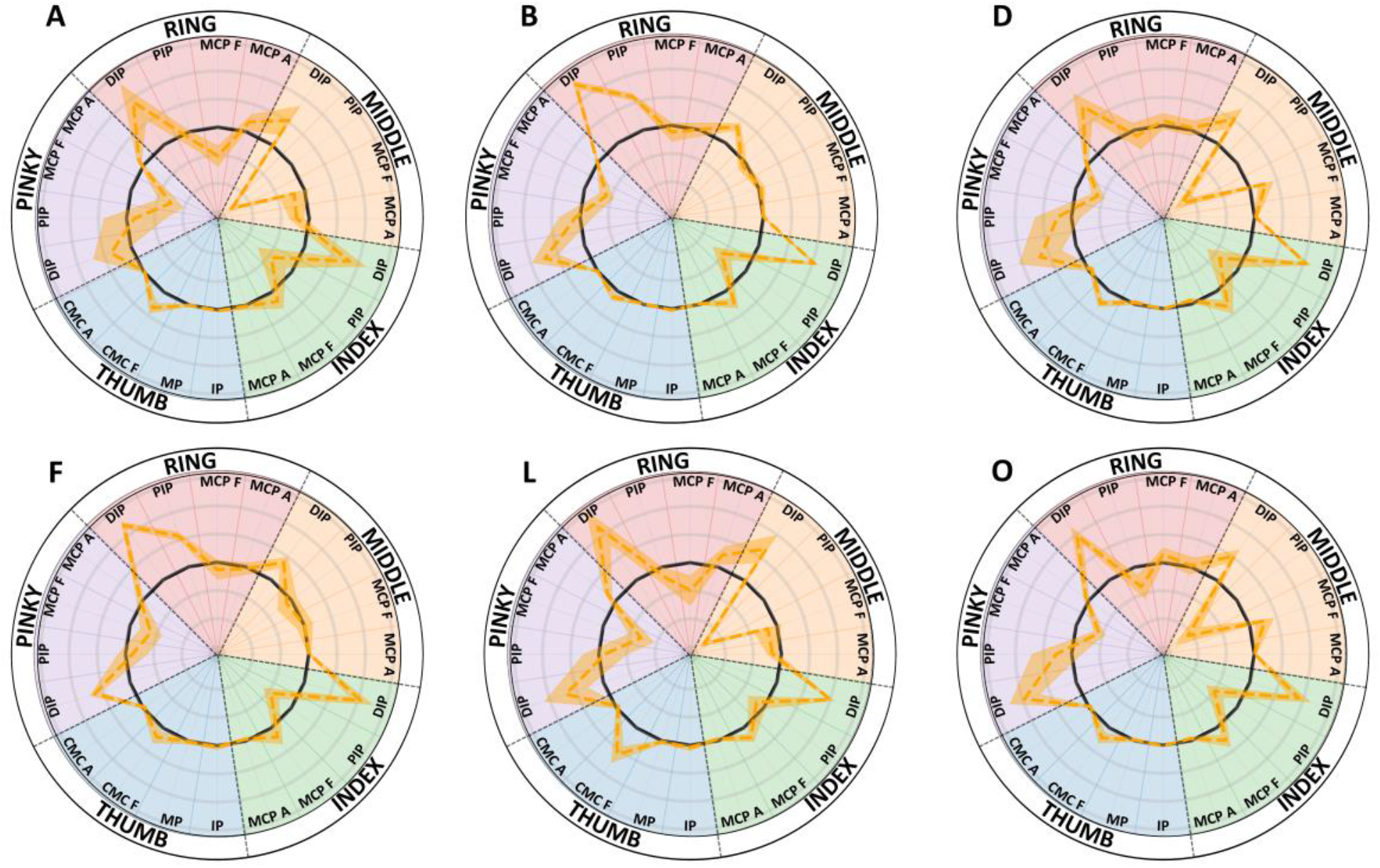
Tracking error across hand and thumb joints for all ASL tasks. The orange dashed line represents the signed mean tracking error for each degree of freedom, averaged across all converged trials and participants; shaded regions indicate ±1 standard deviation. The solid black line denotes perfect tracking (0° error). Positive values (outward from 0°) indicate errors in the flexion or abduction direction depending on the joint, whereas negative values (inward) indicate extension or adduction errors. Background colors group joints by digit and match the color scheme used in Figure 5. Light gray concentric rings indicate 20° radial increments. A consistent outward spike is observed at the DIP joint of each finger, reflecting a flexion bias across tasks, whereas other joints exhibit smaller errors with more variable directional patterns.

## Notes

### Competing Interest Statement

The authors have declared no competing interest.

## References

1. Gordon AM, Malik AT, Goyal KS. Trends of hand injuries presenting to US emergency departments: A 10-year national analysis. The American Journal of Emergency Medicine. 2021;50:466–471. doi:10.1016/j.ajem.2021.08.059

2. Tamulevicius M, Bucher F, Dastagir N, Maerz V, Vogt PM, Dastagir K. Demographic shifts reshaping the landscape of hand trauma: a comprehensive single-center analysis of changing trends in hand injuries from 2007 to 2022. Inj Epidemiol. 2024;11(1):25. doi:10.1186/s40621-024-00510-8

3. How many people are affected by/at risk for stroke? | NICHD - Eunice Kennedy Shriver National Institute of Child Health and Human Development. January 3, 2025. Accessed August 26, 2025. https://www.nichd.nih.gov/health/topics/stroke/conditioninfo/risk

4. Barth J, Geed S, Mitchell A, Lum PS, Edwards DF, Dromerick AW. Characterizing upper extremity motor behavior in the first week after stroke. PLOS ONE. 2020;15(8):e0221668. doi:10.1371/journal.pone.0221668

5. CDC. Data and Statistics for Cerebral Palsy | CDC. Centers for Disease Control and Prevention. December 30, 2020. Accessed August 22, 2025. https://archive.cdc.gov/ncbddd/cp/data.html

6. Makki D, Duodu J, Nixon M. Prevalence and pattern of upper limb involvement in cerebral palsy. J Child Orthop. 2014;8(3):215–219. doi:10.1007/s11832-014-0593-0

7. Moellhoff N, Throner V, Frank K, et al. Epidemiology of hand injuries that presented to a tertiary care facility in Germany: a study including 435 patients. Arch Orthop Trauma Surg. 2023;143(3):1715–1724. doi:10.1007/s00402-022-04617-9

8. Work Injuries and Illnesses by Part of Body. Injury Facts. Accessed August 26, 2025. https://injuryfacts.nsc.org/work/industry-incidence-rates/work-injuries-and-illnesses-by-part-of-body/

9. Abdullah M, Hulleck AA, Katmah R, Khalaf K, El-Rich M. Multibody dynamics-based musculoskeletal modeling for gait analysis: a systematic review. J NeuroEngineering Rehabil. 2024;21(1):178. doi:10.1186/s12984-024-01458-y

10. Current Perspectives on the Biomechanical Modelling of the Human Lower Limb: A Systematic Review | Archives of Computational Methods in Engineering. Accessed August 26, 2025. https://link.springer.com/article/10.1007/s11831-019-09393-1

11. Bolsterlee B, Veeger DHEJ, Chadwick EK. Clinical applications of musculoskeletal modelling for the shoulder and upper limb. Med Biol Eng Comput. 2013;51(9):953–963. doi:10.1007/s11517-013-1099-5

12. Steele KM, Demers MS, Schwartz MH, Delp SL. Compressive tibiofemoral force during crouch gait. Gait Posture. 2012;35(4):556–560. doi:10.1016/j.gaitpost.2011.11.023

13. Mokhtarzadeh H, Yeow CH, Hong Goh JC, Oetomo D, Malekipour F, Lee PVS. Contributions of the soleus and gastrocnemius muscles to the anterior cruciate ligament loading during single-leg landing. J Biomech. 2013;46(11):1913–1920. doi:10.1016/j.jbiomech.2013.04.010

14. Reinbolt JA, Seth A, Delp SL. Simulation of human movement: applications using OpenSim. Procedia IUTAM. 2011;2:186–198. doi:10.1016/j.piutam.2011.04.019

15. Steele KM, Seth A, Hicks JL, Schwartz MS, Delp SL. Muscle contributions to support and progression during single-limb stance in crouch gait. J Biomech. 2010;43(11):2099–2105. doi:10.1016/j.jbiomech.2010.04.003

16. Winby CR, Lloyd DG, Besier TF, Kirk TB. Muscle and external load contribution to knee joint contact loads during normal gait. Journal of Biomechanics. 2009;42(14):2294–2300. doi:10.1016/j.jbiomech.2009.06.019

17. Molinaro DD, Scherpereel KL, Schonhaut EB, Evangelopoulos G, Shepherd MK, Young AJ. Task-agnostic exoskeleton control via biological joint moment estimation. Nature. 2024;635(8038):337–344. doi:10.1038/s41586-024-08157-7

18. Fregly BJ, Reinbolt JA, Rooney KL, Mitchell KH, Chmielewski TL. Design of patient-specific gait modifications for knee osteoarthritis rehabilitation. IEEE Trans Biomed Eng. 2007;54(9):1687–1695. doi:10.1109/tbme.2007.891934

19. Rajagopal A, Kidziński Ł, McGlaughlin AS, Hicks JL, Delp SL, Schwartz MH. Pre-operative gastrocnemius lengths in gait predict outcomes following gastrocnemius lengthening surgery in children with cerebral palsy. PLoS One. 2020;15(6):e0233706. doi:10.1371/journal.pone.0233706

20. Engelhardt L, Melzner M, Havelkova L, et al. A new musculoskeletal AnyBodyTM detailed hand model. Computer Methods in Biomechanics and Biomedical Engineering. 2020;24(7):777–787. doi:10.1080/10255842.2020.1851367

21. Mirakhorlo M, Van Beek N, Wesseling M, Maas H, Veeger HEJ, Jonkers I. A musculoskeletal model of the hand and wrist: model definition and evaluation. Computer Methods in Biomechanics and Biomedical Engineering. 2018;21(9):548–557. doi:10.1080/10255842.2018.1490952

22. McFarland DC, Binder-Markey BI, Nichols JA, Wohlman SJ, de Bruin M, Murray WM. A Musculoskeletal Model of the Hand and Wrist Capable of Simulating Functional Tasks. IEEE Trans Biomed Eng. 2023;70(5):1424–1435. doi:10.1109/TBME.2022.3217722

23. Lee JH, Asakawa DS, Dennerlein JT, Jindrich DL. Finger Muscle Attachments for an OpenSim Upper-Extremity Model. PLOS ONE. 2015;10(4):e0121712. doi:10.1371/journal.pone.0121712

24. Binder-Markey BI, Murray WM. Incorporating the length-dependent passive-force generating muscle properties of the extrinsic finger muscles into a wrist and finger biomechanical musculoskeletal model. Journal of Biomechanics. 2017;61:250–257. doi:10.1016/j.jbiomech.2017.06.026

25. Crouch DL, Huang H. Musculoskeletal model predicts multi-joint wrist and hand movement from limited EMG control signals. In: 2015 37th Annual International Conference of the IEEE Engineering in Medicine and Biology Society (EMBC). 2015:1132–1135. doi:10.1109/EMBC.2015.7318565

26. Zhao Y, Zhang Z, Li Z, Yang Z, Dehghani-Sanij AA, Xie S. An EMG-Driven Musculoskeletal Model for Estimating Continuous Wrist Motion. IEEE Transactions on Neural Systems and Rehabilitation Engineering. 2020;28(12):3113–3120. doi:10.1109/TNSRE.2020.3038051

27. Blana D, Chadwick EK, van den Bogert AJ, Murray WM. Real-time simulation of hand motion for prosthesis control. Computer Methods in Biomechanics and Biomedical Engineering. 2017;20(5):540–549. doi:10.1080/10255842.2016.1255943

28. Melzner M, Engelhardt L, Simon U, Dendorfer S. Electromyography-Based Validation of a Musculoskeletal Hand Model. J Biomech Eng. 2021;144(021005). doi:10.1115/1.4052115

29. Vignais N, Marin F. Analysis of the musculoskeletal system of the hand and forearm during a cylinder grasping task. International Journal of Industrial Ergonomics. 2014;44(4):535–543. doi:10.1016/j.ergon.2014.03.006

30. Improved Trajectory Reconstruction for Markerless Pose Estimation | IEEE Conference Publication | IEEE Xplore. Accessed August 27, 2025. https://ieeexplore.ieee.org/abstract/document/10340745

31. Biomechanical Arm and Hand Tracking with Multiview Markerless Motion Capture | IEEE Conference Publication | IEEE Xplore. Accessed August 27, 2025. https://ieeexplore.ieee.org/document/10719940

32. OpenSim Moco: Musculoskeletal optimal control | PLOS Computational Biology. Accessed August 27, 2025. https://journals.plos.org/ploscompbiol/article?id=10.1371/journal.pcbi.1008493

33. Millard M, Uchida T, Seth A, Delp SL. Flexing Computational Muscle: Modeling and Simulation of Musculotendon Dynamics. J Biomech Eng. 2013;135(021005). doi:10.1115/1.4023390

34. Evaluation of Direct Collocation Optimal Control Problem Formulations for Solving the Muscle Redundancy Problem | Annals of Biomedical Engineering. Accessed August 27, 2025. https://link.springer.com/article/10.1007/s10439-016-1591-9

35. Saul KR, Hu X, Goehler CM, et al. Benchmarking of dynamic simulation predictions in two software platforms using an upper limb musculoskeletal model. Comput Methods Biomech Biomed Engin. 2015;18(13):1445–1458. doi:10.1080/10255842.2014.916698

36. Practical Methods for Optimal Control and Estimation Using Nonlinear Programming, Second Edition | SIAM Publications Library. Advances in Design and Control. Accessed January 30, 2026. https://epubs.siam.org/doi/book/10.1137/1.9780898718577

37. Wächter A, Biegler LT. On the implementation of an interior-point filter line-search algorithm for large-scale nonlinear programming. Math Program. 2006;106(1):25–57. doi:10.1007/s10107-004-0559-y

38. Delp SL, Anderson FC, Arnold AS, et al. OpenSim: Open-Source Software to Create and Analyze Dynamic Simulations of Movement. IEEE Transactions on Biomedical Engineering. 2007;54(11):1940–1950. doi:10.1109/TBME.2007.901024

39. Hicks JL, Uchida TK, Seth A, Rajagopal A, Delp SL. Is My Model Good Enough? Best Practices for Verification and Validation of Musculoskeletal Models and Simulations of Movement. J Biomech Eng. 2015;137(2):0209051–02090524. doi:10.1115/1.4029304

40. Lee SW, Chen H, Towles JD, Kamper DG. Effect of Finger Posture on the Tendon Force Distribution Within the Finger Extensor Mechanism. J Biomech Eng. 2008;130(5):051014. doi:10.1115/1.2978983

41. El-Shennawy M, Nakamura K, Patterson RM, Viegas SF. Three-dimensional kinematic analysis of the second through fifth carpometacarpal joints. J Hand Surg Am. 2001;26(6):1030–1035. doi:10.1053/jhsu.2001.28761

42. Kuhns JG. Structural and Dynamic Bases of Hand Surgery. JAMA. 1969;207(8):1521. doi:10.1001/jama.1969.03150210105028

43. Wei Y, Zou Z, Qian Z, Ren L, Wei G. Biomechanical Analysis of the Effect of the Finger Extensor Mechanism on Hand Grasping Performance. IEEE Transactions on Neural Systems and Rehabilitation Engineering. 2022;30:360–368. doi:10.1109/TNSRE.2022.3146906

44. Jadelis C, Ellis B, Kamper D, Saul KR. Cosimulation of the Index Finger Extensor Apparatus with Finite Element and Musculoskeletal Models. J Biomech. 2023;157:111725. doi:10.1016/j.jbiomech.2023.111725

45. Dogadov AA, Valero-Cuevas FJ, Servière C, Quaine F. The Bundles of Intercrossing Fibers of the Extensor Mechanism of the Fingers Greatly Influence the Transmission of Muscle Forces. International Journal for Numerical Methods in Biomedical Engineering. 2025;41(7):e70068. doi:10.1002/cnm.70068

